# Transcriptome profiling of the branchial arches reveals cell type composition and a conserved signature of neural crest cell invasion

**DOI:** 10.1101/2020.02.28.969915

**Authors:** Jason A Morrison, Rebecca McLennan, Jessica M Teddy, Allison R Scott, Jennifer C Kasemeier-Kulesa, Madelaine M Gogol, Paul M Kulesa

## Abstract

The vertebrate branchial arches that give rise to structures of the head, neck, and heart form with very dynamic tissue growth and well-choreographed neural crest, ectoderm, and mesoderm cell dynamics. Although this morphogenesis has been studied by marker expression and fate-mapping, the mechanisms that control the collective migration and diversity of the neural crest and surrounding tissues remain unclear, in part due to the effects of averaging and need for cell isolation in conventional transcriptome analysis experiments of multiple cell populations. We used label free single cell RNA sequencing on 95,000 individual cells at 2 developmental stages encompassing formation of the first four chick branchial arches to measure the transcriptional states that define the cellular hierarchy and invasion signature of the migrating neural crest. The results confirmed basic features of cell type diversity and led to the discovery of many novel markers that discriminate between axial level and distal-to-proximal cell populations within the branchial arches and neural crest streams. We identified the transcriptional signature of the most invasive neural crest that is conserved within each branchial arch stream and elucidated a set of genes common to other cell invasion signatures in types in cancer, wound healing and development. These data robustly delineate molecularly distinct cell types within the branchial arches and identify important molecular transitions within the migrating neural crest during development.

## INTRODUCTION

During vertebrate development, multipotent neural crest cells migrate in discrete streams from the dorsal neural tube to the periphery to form functional structures (Etchevers 2019). Mistakes in collective migration or cell differentiation of the neural crest can result in severe birth defects (Frisdal and Trainor 2014). Therefore, to understand how a healthy individual develops, it is important that we understand how this coordinated movement of cells is controlled. Moreover, the neural crest is a well-known in vivo model to test hypothetical mechanisms described in other cell invasion phenomena, such as cancer and wound healing, since migrating cells are accessible to time-lapse imaging, molecular perturbation and profiling (Kulesa 2013a,b). Thus, knowledge of the cellular and molecular mechanisms underlying neural crest cell migration will lead to a better understanding of structural birth defects and insights into the hallmarks of collective cell motion that is prevalent throughout development and disease.

Neural crest cell migration begins at the midbrain and continues in a rostral-to-caudal manner all along the vertebrate axis. In the hindbrain, neural crest cells that emerge from rhombomere 1 (r1) through r7 are shaped into discrete migratory streams that invade the first four branchial arches (BA1-4). These branchial arches are sculpted into important structures in the face, neck and heart (Frisdal and Trainor, 2014). Characterization of neural crest migration from time-lapse imaging data has revealed a rich set of cell behaviors that include cell contact guidance, contact inhibition of locomotion, chemotaxis and cell communication that together promote collective cell movement (Giniunaite 2019a; Giniunaite 2019b; Szabó and Mayor 2018). Signals within the dorsal neural tube and microenvironments through which cells travel are coordinated in space and time to control cell exit locations, prevent stream mixing, and direct cells into the correct branchial arch (Kulesa and McLennan, 2015). Despite the wealth of cell behavioral data, much less is known about the network of genes that underlie this complex and well-choreographed cellular hierarchy and collective migration of the neural crest.

To begin to address these questions, we previously conducted single cell analysis of cranial neural crest cells migrating into the second branchial arch (BA2) at three progressive developmental stages: delamination from the dorsal neural tube, migration and BA2 invasion. We showed that gene expression is dynamic within the neural crest cell migratory stream and depends on cell position and time after exit from the neural tube (McLennan 2012; McLennan 2015a,b; Morrison 2017a). We identified gene expression differences in leader versus follower neural crest cells, including a consistent transcriptional signature associated with a subset of cells at the invasive front, which we termed ‘Trailblazers’ (Morrison 2017a). Since discrete neural crest cell migratory streams populate the first four branchial arches and form distinct structures, it is important to understand the degree of similarity and differences in the molecular heterogeneity within the first four branchial arch streams. Further, it is unclear how signals in the branchial arch microenvironments are coordinated to drive collective migration and cell type diversity.

In this study, we take advantage of methods that permit label free single cell RNA sequencing (scRNA-seq) to efficiently profile populations of individual cells in complex tissues. Specifically, we measure transcriptional states of migrating neural crest cells and surrounding tissues from BA1-BA4 of chicken embryos at 2 developmental stages. To guide bioinformatics analysis and aid in the identification of neural crest and other cell types, we utilized our previously generated gene expression database of the BA2 neural crest cell migratory stream as a reference (McLennan 2012; McLennan 2015a,b; Morrison 2017a). This approach enabled us to determine spatio-temporal gene expression profiles and compare differences in front (distal 20%) and back (proximal 80%) cell subpopulations within all four branchial arches. We uncovered profiles associated with the molecular transition to the most invasive neural crest cells and identification a distinct transcriptional signature of these cells. An integrated multiplexed fluorescence in-situ hybridization strategy (Morrison 2017b) enabled us to examine a subset of Trailblazer genes, validate the analyses and investigate changes in in vitro neural crest cell behaviors after loss-of-function of a small subset of Trailblazer genes. Lastly, through comparisons with published gene expression signatures from a wide range of other cell invasion phenomena our analyses identify a subset of genes shared with the neural crest Trailblazer signature. These results represent a comprehensive analysis of the cellular hierarchy and molecular heterogeneity of the migrating neural crest and BA1-BA4 tissues during vertebrate development.

## RESULTS

### Label free and unsorted scRNA-seq identified distinct cell types within BA1-BA4

We previously demonstrated that cranial neural crest cells that exit from the mid-hindbrain and travel to BA2 express a subset of genes that depend on cell position within the migratory stream and the timing of their emergence from the dorsal neural tube (Morrison 2017a). Neural crest cells within the front 20% (distal) subpopulation have a distinct transcriptional signature in comparison to the back 80% (proximal) neural crest cell subpopulation. These distinct molecular signatures vary in time as cells travel through different microenvironments to invade BA2. We also found that a subset of the invasive cells at the leading edge, termed ‘Trailblazers’, express a unique and consistent transcriptional signature throughout the migratory period. It was unknown whether these spatio-temporal molecular heterogeneities and the unique Trailblazer transcriptional signature are also present within neighboring BA1, BA3, and BA4 neural crest cell migratory streams and what microenvironmental signals may underlie these profiles.

To address these questions, we expanded the single cell transcriptome analysis along the anterior-posterior axis to investigate patterns of expression within BA1-4 (Fig. 1A-B). We dissected chick BA1 and BA2 at Hamburger and Hamilton (HH)13 and BA3 and BA4 at HH15 which represent developmental timepoints during neural crest cell migration and branchial arch formation (Hamburger and Hamilton 1951). Each branchial arch microenvironment was subdivided into the front and the back (Fig. 1A-B). scRNA-seq (10x Genomics Chromium) and bioinformatic analyses (Seurat) of label free and unsorted branchial arch tissues produced approximately 95,000 single cell transcriptional profiles, containing cell types such as ectoderm, mesoderm, and migrating neural crest cells (Butler 2018) (Fig. 1B and Suppl. Fig. 1). We next utilized Uniform Manifold Approximation and Projection (UMAP) analysis to display the transcriptomes as 7 distinct clusters, with each of the clusters marked by increased expression of unique genes (Becht 2019; McInnes 018) (Fig. 1C-F; Suppl. Table 1).

**Figure 1.**
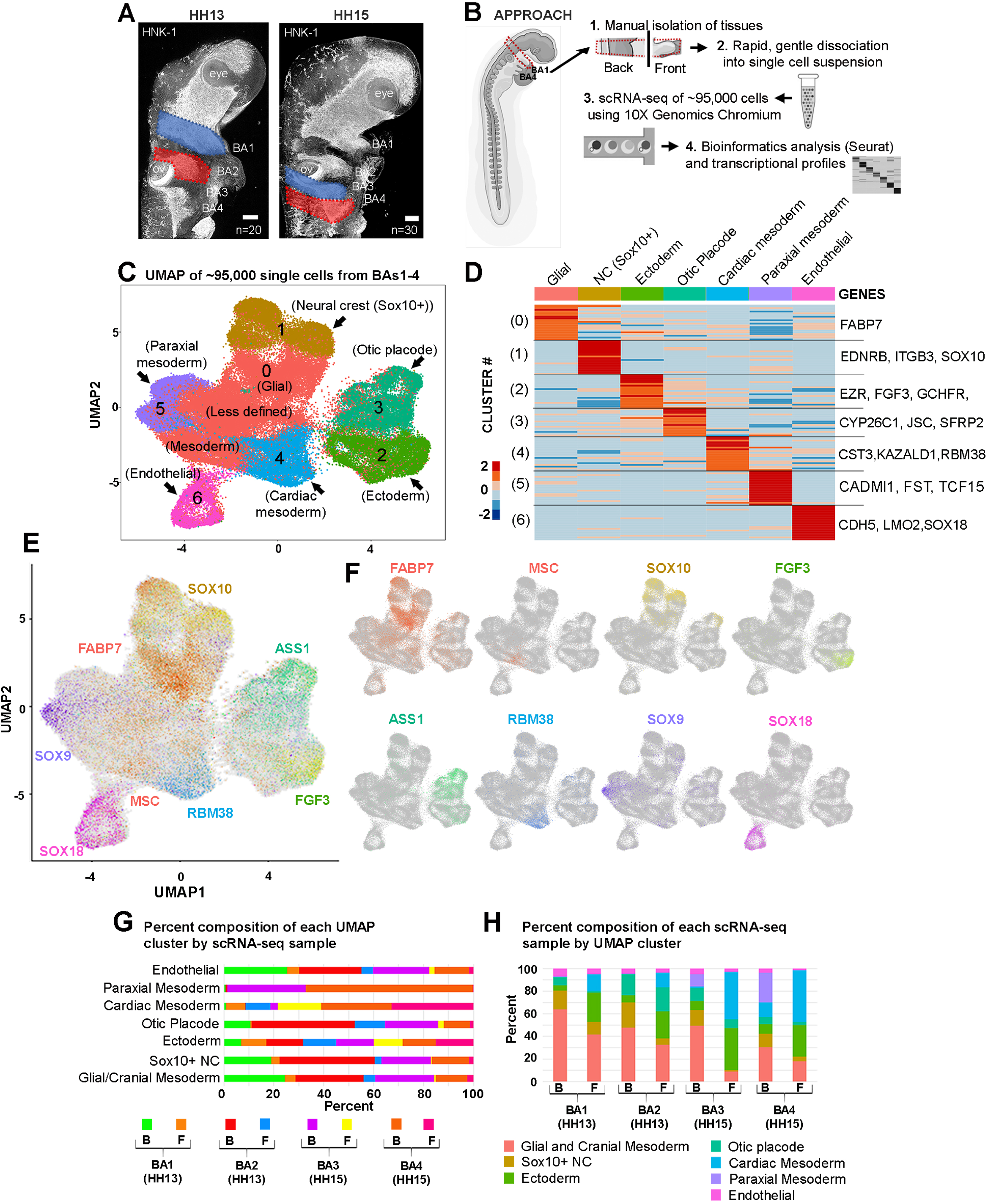
Single cell sequencing identifies distinct cell types within embryonic branchial arches. (A) Unperturbed chick branchial arches (BA) 1-4 harvested for single cell RNA-seq analysis during active neural crest migration. (B) Isolation, scRNA-seq and bioinformatic analysis of BA1-4 (C) UMAP segregates ∼95,000 single cell transcriptomes from BA1-4 into 7 clusters. (D) 7 clusters of BA1-4 cells are distinguished by upregulation of marker genes. (E-F) UMAP of all BA1-4 single cell transcriptomes color-coded by expression of individual genes that mark specific clusters. (G) Contribution of each spatially segregated sample to each of the 7 UMAP clusters. (H) Cell type composition of each of the spatially segregated scRNA-seq samples. Branchial Arch (BA); Hamburger & Hamilton chick developmental stage (HH); Neural Crest (NC); Front (F); Back (B).

In order to more fully characterize these 7 clusters, including the spatial location of the clusters within the context of the developing embryo conferred by the 8 spatially unique samples (Fig. 1A,B), we determined the contribution of each sample to each cluster (Fig. 1G, Suppl. Table 2). The paraxial mesoderm cluster is dominated by cells from the back of BA3-4 and most of the cardiac mesoderm cluster is made up of cells from BA3-4 (Fig. 1G). Contributions to endothelial, ectodermal, SOX10-positive neural crest, glial and mesodermal cells are observed from all branchial arches analyzed.

To better understand the types of cells and tissues that exist at each anatomical location within BA1-4, we calculated the percentage composition of each of the 8 scRNA-seq samples (Fig. 1H). The front of each branchial arch contained a higher percentage of ectodermal cells than the back and conversely, the back of each branchial arch had a higher percentage of glia and cranial mesoderm than the front. The percentage of endothelial cells was consistent across all samples. As noted in Fig. 1G, paraxial mesoderm was almost exclusive to the back samples from BA3-4 and cardiac mesoderm was preferentially found in cells from BA3-4 (Fig. 1H).

### Expression of segmental HOX genes validates scRNA-seq of cranial to cardiac branchial arches

To further characterize the scRNA-seq analysis of BA1-4 tissue, we examined the axially-restricted profiles of expression of Homeobox (HOX; HOXA2, HOXA3, HOXB3), Distal-less Homeobox 5 (DLX5), and MAF BZIP Transcription Factor B (MAFB) genes (Suppl. Fig. 2A-F). When isolating BA1-4 tissue for scRNA-seq, both axial level and proximal-distal spatial information was maintained within individual samples (Fig. 1A-B). The SOX10-positive neural crest cells (cluster 1) segregates into 3 distinct subclusters corresponding to the branchial arches that they invade (Suppl. Fig. 2A). To evaluate expression of known arch-specific HOX genes within cells from individual arches, we plotted the expression of DLX5, individual HOX genes and MAFB within the context of our UMAP (Suppl. Fig. 2B-F). DLX5 is expressed in all subclusters of SOX10-positive neural crest cells (Suppl. Fig. 2E), supporting previous bulk RNA-seq data (Simoes-Costa 2014). The SOX10-positive neural crest cell subcluster, composed of cells from BA1, is devoid of HOX expression, while. HOXA2 is specifically enriched in BA2 cells and the expression of HOXA3, HOXB3, and MAFB are restricted to BA3-4 (Suppl. Fig. 2A-F). These data confirm the appropriate localized expression of genes with well-characterized segmentation patterns within the hindbrain and validate the effectiveness of the dissection and isolation protocol.

### Neural crest cell type composition is determined by axial level and proximal-to-distal position within each branchial arch

To better understand similarities and differences along the axis within BA1-4, we highlighted pre-otic and post-otic cells within the context of the 7 cluster UMAP (Fig. 2). We visualized the contribution of each spatially-restricted sample to the entirety of the UMAP (Fig. 2D-I) and find that both BA1-2 and BA3-4 cells supply ectoderm, otic placode, cardiac mesoderm and endothelial clusters. For example, endothelial cells are equally distributed among pre-and post-otic regions (Fig. 1G-H, 2A-C,E,H). Conversely, cells within SOX10-positive neural crest, cardiac mesoderm and paraxial mesoderm clusters segregate based upon their pre-otic and post-otic origin (compare differences in black and magenta regions in Fig. 2A-C). Cranial mesoderm marked by Cytochrome P450 Family 26 Subfamily C Member 1 (CYP26C1) showed a dominant contribution from pre-otic cells as previously observed (Bothe 2011). Meis Homeobox 2 (MEIS2) expression is specific to post-otic cells, especially in SOX10-positive neural crest and paraxial mesoderm (Fig. 2A-H).

**Figure 2.**
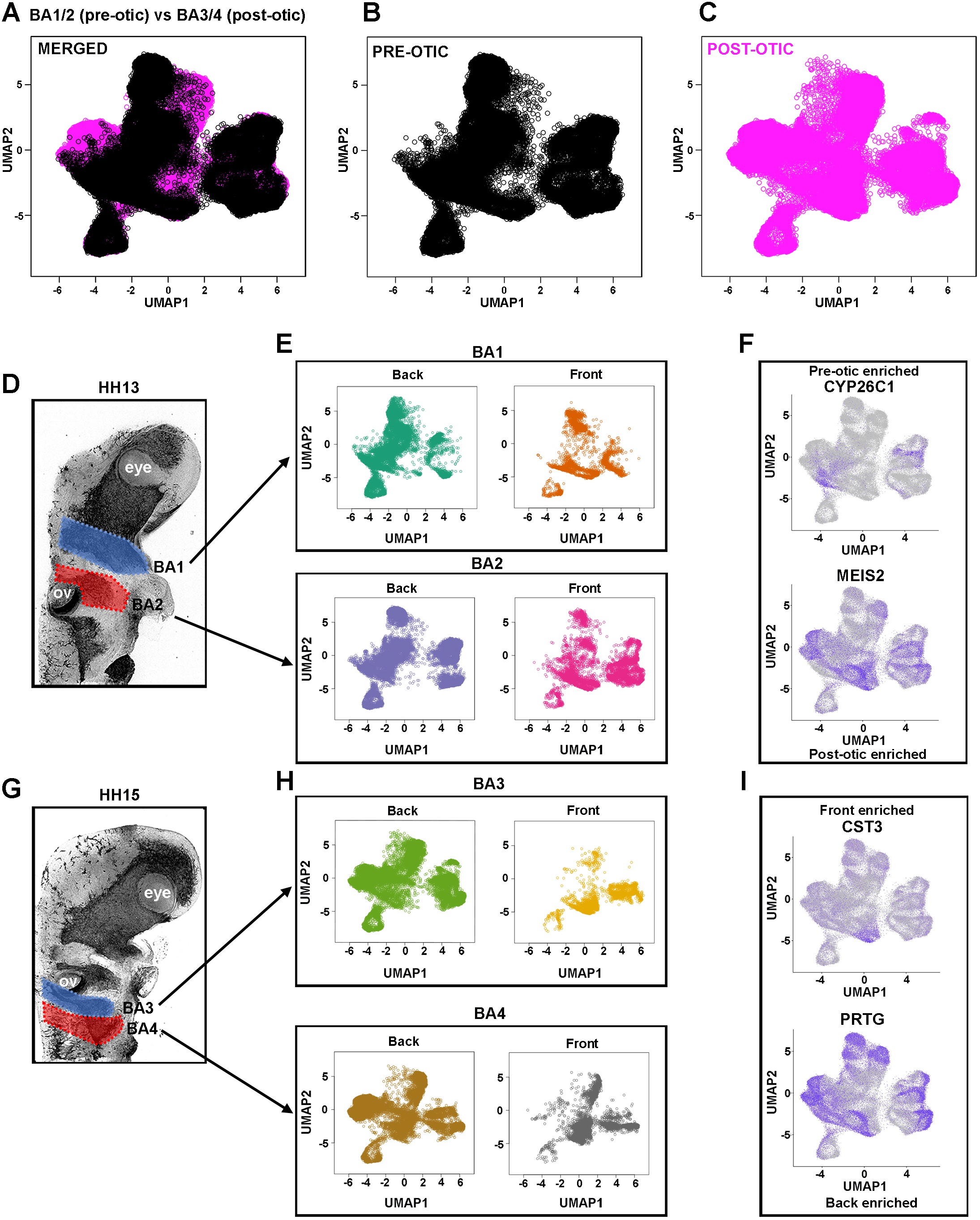
Cellular composition is determined by axial level and position within each branchial arch (proximal vs distal). (A-C) UMAP of BA1-4 single cell transcriptomes color-coded by pre-otic (black) or post-otic (magenta) location. (D) Unperturbed HH13 branchial arches 1-2 harvested for single cell RNA-seq analysis. (E) UMAP of BA1-4 single cell transcriptomes color-coded by contribution of HH13 BA1-2 Front and Back samples. (F) UMAP of BA1-4 single cell transcriptomes color-coded by expression of genes enriched in pre-otic (CYP26C1) and post-otic (MEIS2) cells. (G) Unperturbed HH15 branchial arches 3-4 harvested for single cell RNA-seq analysis. (H) UMAP of BA1-4 single cell transcriptomes color-coded by contribution of HH15 BA3-4 Front and Back samples. (I) UMAP of BA1-4 transcriptomes color-coded by expression of genes enriched in Front (CST3) and Back (PRTG) cells. Branchial Arch (BA); Hamburger & Hamilton chick developmental stage (HH).

In a previous study, we identified distinct leader and follower neural crest molecular profiles within the BA2 neural crest cell migratory stream (Morrison 2017a). For comparison, we distinguished proximal and distal subregions of the arches within the context of the 7 cluster UMAP. There is a significant overlap between the front and back cells in ectoderm and endothelial clusters (Fig. 2E,H) suggesting a similar transcriptional signature for front and back cells within these clusters. In contrast, front and back cells displayed some spatial segregation within the cluster of SOX10-positive neural crest cells at each branchial arch level (Fig. 2E,H). Other clusters were also comprised of disproportionate numbers of front and back cells. For example, cardiac mesoderm showed a higher proportion of front cells and otic placode and paraxial mesoderm are primarily made up of cells at the back. Both of these results confirm known biological observations and further validate our scRNA-seq data. We next identified individual genes whose expression differed between front and back (Fig. 2I). CST3 preferentially marks front cells of the cardiac mesoderm (cluster 4) and expression of the neural marker Protogenin (PRTG) is restricted to back cells within most clusters (Fig. 2I). These data suggest that gene expression is highly similar in some cell types (for example, ectoderm and endothelial) despite axial level and proximal-distal position. However other cell types, such as neural crest and mesoderm, show differential expression based upon axial level and/or proximal-distal position, in agreement with our previous analyses of BA2.

### Differences in SOX10 and other traditional markers in the most invasive cells

Migrating neural crest cells are traditionally identified by the expression of neural crest genes, such as SOX10 or ITGB3 (Tucker 1984; Vincent 1983; Southard-Smith 1998; Pietri 2003). Hence, we examined their expression in clusters within the UMAP to identify which contained migrating neural crest cells. We observed expression of SOX10 in neural crest, glia and the otic placode (Fig. 1E-F; clusters 0,1,3). To confirm expression of SOX10 in the migrating neural crest cells and otic vesicle, we used multiplexed fluorescence in situ hybridization combined with immunohistochemistry for HNK-1 (neural crest-specific membrane marker) at two developmental time points (Fig. 3A-D) (Morrison 2017b). There is a striking reduction of SOX10 at the invasive front of all four neural crest cell migratory streams (Fig. 3A-D). Higher magnification of the BA2 neural crest migratory stream at HH13 and HH15 distinctly illustrate this difference, as seen by comparing arrows denoting the distal-most HNK1-positive neural crest (turquoise) with asterisks marking a dramatic reduction in SOX10 expression (yellow) (Fig. 3B,D). Novel markers of migrating neural crest cells were identified and included: Ubiquitin Like Modifier Acting Enzyme 7 (UBA7), Inositol-Tetrakisphosphate 1-Kinase (ITPK1), and Collagen Type XVIII Alpha 1 Chain (COL18A1) (Suppl. Fig. 2H). These results strengthen our previous observation that SOX10 expression appeared to be reduced as neural crest cells migrated further from the dorsal neural tube exit location and invaded BA2 (McLennan 2012; McLennan 2015a,b; Morrison 2017a,b). Furthermore, they provide a new subset of markers for exploring temporal dynamic signatures in migrating neural crest cells.

**Figure 3.**
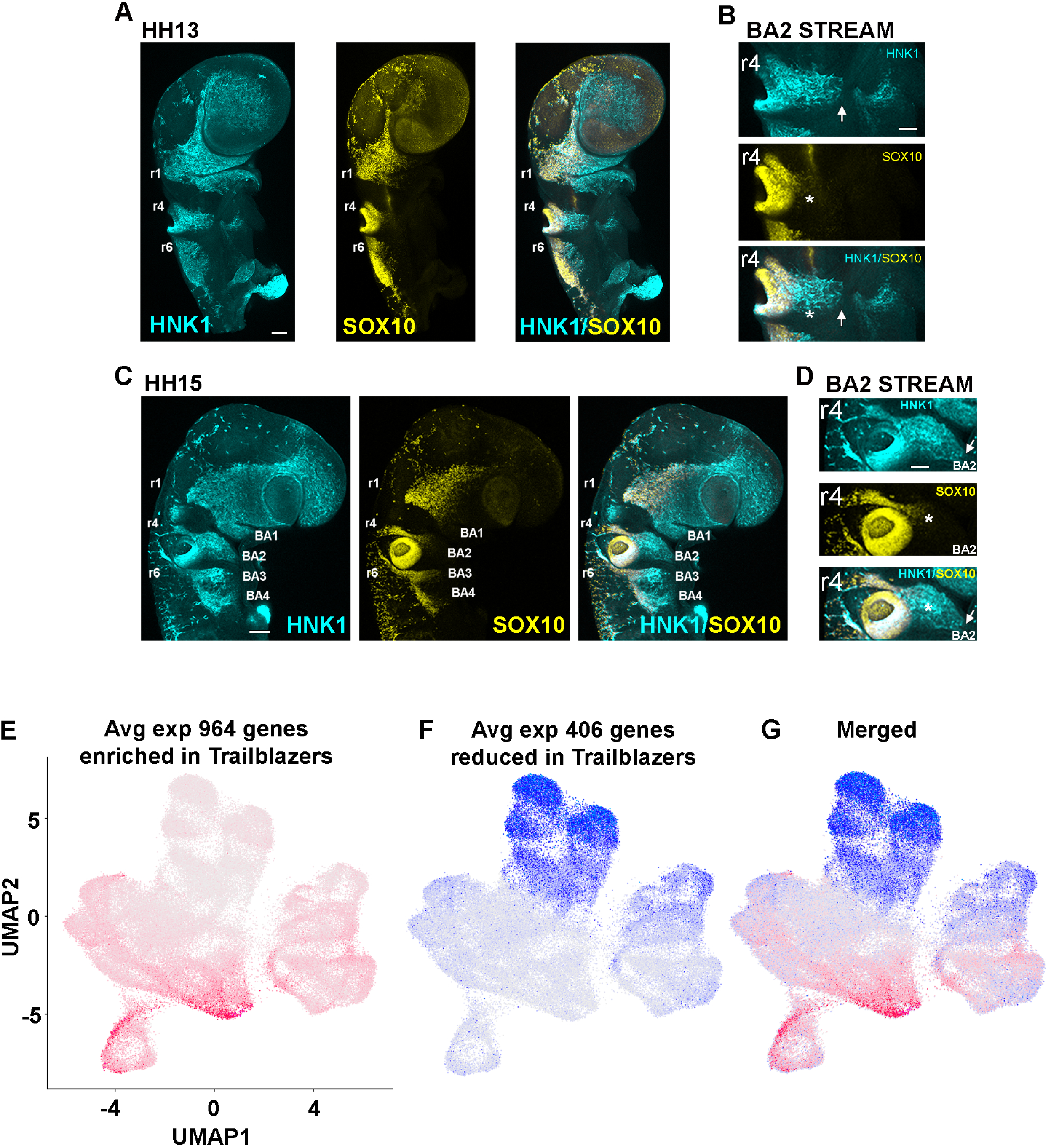
SOX10+ neural crest cells contribute to multiple cell types and cells enriched for the Trailblazer neural crest profile exist at the distal portion of each branchial arch. (A-D) At HH13 and 15, migrating all cranial-to-cardiac neural crest cells are strongly labeled by HNK-1 (arrows), while SOX10 expression (asterisks) inefficiently labels neural crest cells at the migratory front. (E) UMAP of BA1-4 single cell transcriptomes color-coded by the average expression of 964 genes enriched in BA2 Trailblazer NC cells compared to all other BA2 NC. (F) UMAP of BA1-4 single cell transcriptomes color-coded by the average expression of 406 genes reduced in BA2 Trailblazer NC cells compared to all other BA2 NC. (G) Merged images of E and F. Branchial Arch (BA); Hamburger & Hamilton chick developmental stage (HH); Rhombomeres 1-6 (r1-6). Scalebars are 100um (A,C) and 50um (B,D).

### The Trailblazer transcriptional signature is conserved within BA1-4 neural crest cell migratory streams

Next, we explored whether the novel Trailblazer transcriptional signature associated with the most invasive neural crest cells migrating into BA2 is conserved in other branchial arches (Morrison 2017a). Since SOX10 and other commonly used neural crest markers are reduced in Trailblazers (Fig. 3A-D), we asked whether Trailblazers could be more accurately identified within the UMAP of BA1-4 tissue by employing the known BA2 Trailblazer molecular profile as a reference. The cardiac mesoderm and endothelial clusters (Fig. 1C; clusters 4 and 6, respectively) were enriched for an average of the 964 genes enriched in the BA2 Trailblazer signature (Fig. 3E). As cluster 6 distinctly expressed well-characterized markers of endothelial cells, such as SOX18, LMO2 and CDH5 (Fig. 1C-F; Fig. 3E; Suppl. Table 1), we instead focused our efforts to identify Trailblazer neural crest cells within the cardiac mesoderm cluster. Since the BA2 Trailblazer signature was originally determined by comparing Trailblazer cells within BA2 to the other neural crest cell streams, enrichment of BA2 Trailblazer genes in the cardiac mesoderm cluster should be compared to expression within the SOX10-positive neural crest cluster (Fig. 1C; cluster 1). We observe that unlike the cardiac mesoderm cluster, the SOX10-positive neural crest cluster is not enriched for trailblazer genes, suggesting it may correspond to a more proximal population of neural crest cells (Fig. 3E). To further verify similarity of the cardiac mesoderm and BA2 Trailblazer molecular profiles, we looked at the average expression of the 406 genes reduced in the Trailblazer signature (Fig. 3F). There is a clear reduction of genes reduced in Trailblazers in the cardiac mesoderm cluster compared to the SOX10-positive neural crest cluster 1 (Fig. 1C; cluster 4 and Fig. 3G). These results suggest that Trailblazer neural crest cells are not part of the SOX10+ neural crest cluster (cluster 1), which appear to represent more proximal neural crest cells.

### Trailblazer neural crest cells exist at all axial levels analyzed

The Trailblazer subpopulation of BA2 was a small percentage (∼2%) of cells within the neural crest stream and confined to the invasive front (Morrison 2017a). Here, our analysis of all BA1-4 tissue found a larger percentage (8%) of the cells that clustered as cardiac mesoderm (Suppl. Table 2). The cardiac mesoderm cluster (cluster 4) also contained cells from the back (proximal) portion of BA3-4 (Fig. 1G-H, Suppl. Table 2). These observations suggested that cluster 4 may contain both Trailblazer neural crest in addition to mesodermal cells. To explore the molecular heterogeneity within the cardiac mesoderm cluster, we independently clustered only the cardiac mesoderm cells, and color-coded the resulting single cell transcriptomes by sample types that confer both spatial and temporal information (Fig. 4C, Suppl. Fig. 4). We find that single cell transcriptomes on the left side of the subclustering composition UMAP for cluster 4 are almost exclusively comprised of front and back BA4 cells (Fig. 4C, Suppl. Fig. 4). Single cell transcriptomes on the right side of the plot contained front cells from all four branchial arches and very few back cells.

**Figure 4.**
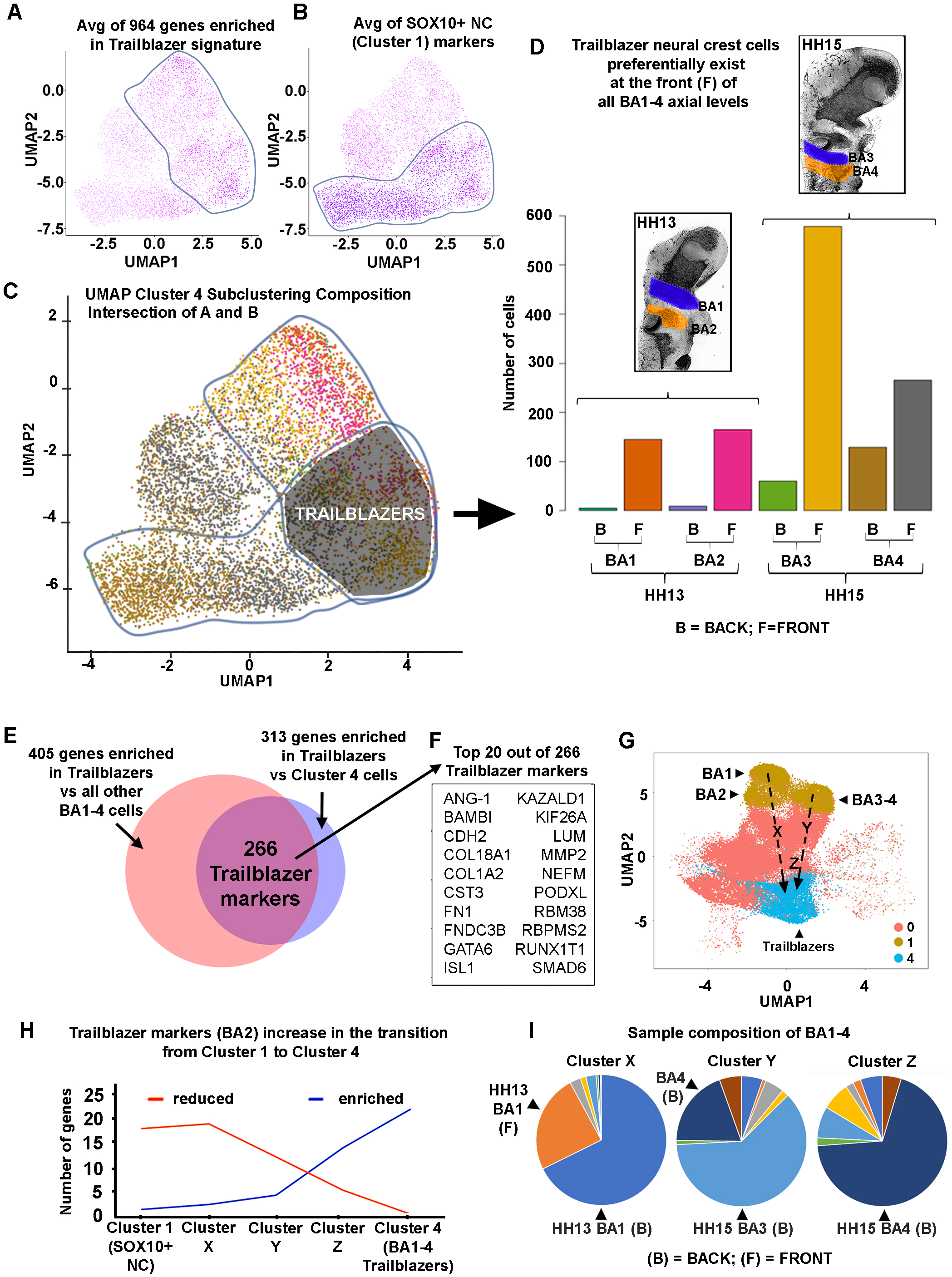
Trailblazer neural crest signature refined by comparison with all branchial arch cells at multiple axial levels. (A) A subset of cardiac mesoderm (cluster 4) cells is enriched for the average expression of 964 genes enriched in BA2 Trailblazer NC cells compared to all other BA2 NC. (B) A subset of cardiac mesoderm (cluster 4) cells is enriched for the average expression of SOX10+ neural crest markers. (C) The intersection of cells enriched for both BA2 Trailblazer and SOX10+ neural crest markers define BA1-4 Trailblazer NC cells. Cells of the cardiac mesoderm (cluster 4) are color-coded by sample type (developmental stage, branchial arch and position within the arch). (D) BA1-4 Trailblazer NC cells preferentially exist at the front of all branchial arches analyzed. (E) 266 markers of BA1-4 Trailblazers were found at the intersection of 405 genes enriched in BA1-4 Trailblazers compared to all other BA1-4 cells and 313 genes enriched in BA1-4 Trailblazers compared to all other cells in the cardiac mesoderm cluster. (F) Twenty BA1-4 Trailblazers compared to all other BA1-4 cells that are also enriched in BA1-4 Trailblazer NC cell markers. (G) UMAP of BA1-4 single cell transcriptomes showingSOX10-positve neural crest (cluster 1), Trailblazer neural crest (subset of cluster 4) and distinct subclusters between clusters 1 and 4 (X, Y and Z). (H) BA2 Trailblazer markers are enriched (blue) in the transition from SOX10+ neural crest to BA1-4 Trailblazer neural crest cells. Conversely, BA2 Trailblazer markers are reduced (orange) in the transition from SOX10+ neural crest to BA1-4 Trailblazer NC cells. (I) Sample composition of BA1-4 clusters containing neural crest cells. Branchial Arch (BA); Hamburger & Hamilton chick developmental stage (HH). Front (F); Back (B).

To determine if the cardiac mesoderm contained Trailblazer neural crest cells, we first examined the expression of the BA2 Trailblazer signature, which contained 964 genes enriched in Trailblazer neural crest compared to other neural crest (Morrison 2017a). We find the average of specifically enriched BA2 Trailblazer markers in single cell transcriptomes on the right side of the cardiac mesoderm UMAP (Fig. 4A; circled), while the bottom portion of cardiac mesoderm cluster is enriched for the average of all SOX10-positive neural crest cell markers (Fig. 4B; circled). We utilized the overlap between of enriched BA2 Trailblazer and neural crest cell markers with the cardiac mesoderm cluster to assign a subpopulation as BA1-4 Trailblazer neural crest cells (Fig. 4C; intersection of circled subregions). To determine whether putative BA1-4 Trailblazers are present at each axial level and if they exist in the more proximal or distal portions of each neural crest cell migratory stream, we utilized the spatial information preserved within the scRNA-seq data set. We find that BA1-4 Trailblazers are almost exclusively at the front of BA1-3 and preferentially at the front of BA4 (Fig. 4D). This analysis of BA1-4 Trailblazer neural crest cells approximated the same Front-Back ratio previously observed for BA2 Trailblazers (Morrison 2017a).

Having identified BA1-4 Trailblazer neural crest cells, we sought to establish their molecular properties. To identify genes unique to BA1-4 Trailblazers, we first compared BA1-4 Trailblazers to all other BA1-4 cells analyzed by scRNA-seq and found 405 genes enriched in the BA1-4 Trailblazer neural crest cells (Fig. 4E; pink circle). Next, we compared BA1-4 Trailblazer neural crest to the transcriptionally similar cardiac mesoderm cells and identified 313 genes enriched in BA1-4 Trailblazer neural crest cells (Fig. 4E; blue circle). At the intersection of these two gene lists are 266 definitive BA1-4 Trailblazer neural crest cell markers (Fig. 4E; magenta circle, Suppl. Table 3) and the top 20 out of 266 genes shown in Fig. 4F). This analysis clearly demonstrates that there is a shared Trailblazer signature for the most invasive neural crest cells in all four branchial arches, indicating that is a conserved property of migrating cranial neural crest streams.

### SOX10-positive to Trailblazer neural crest trajectories contain intermediate subpopulations with reduced multipotency (SOX10) and increased Trailblazer markers

In order to more clearly understand the timing of changes in multipotency within migrating neural crest cells, we focused on changes in SOX10 expression. The scRNA-seq analysis of BA2 neural crest cells showed two distinct transcriptional profiles of recently emigrated cells harvested at HH11, corresponding to recent neural tube exit and acquisition of directed migration (Morrison 2017a). When we closely examined our results at HH11, we observed little or no SOX10 expression in cells with a ‘premigratory’ neural crest cell signature, however, SOX10 was always part of the directed migration expression signature (Suppl. Fig. 3; HH11 cluster 2 (blue bars) neural crest cells). Analysis of BA1-4 cells determined that traditional markers of migrating neural crest, including SOX10, LMO4, EDNRB, ITGB3 and Transcription Factor AP-2 Beta (TFAP2B), were enriched in cluster 1 (Fig. 1D-F; Suppl. Table 1) and to a lesser extent in the glial cluster marked by FABP7 (Fig. 1E-F).

The above results led us to further examine the reduction of multipotency markers in cranial-to-cardiac neural crest cells by analyzing the markers of subpopulations of cells along the axis of SOX10 expression (Fig. 4G; subpopulations X, Y and Z). As traditional neural crest cell markers are downregulated, markers associated with the BA2 Trailblazer neural crest cells emerge (Fig. 4H; compare orange and blue lines plotted from Cluster 1 to Cluster 4). We then analyzed the composition of subclusters X, Y and Z. Strikingly, more than 90% of subcluster X comes from the front and back of BA1 (Fig. 4I). About 80% of the cells in subcluster Y come from the back of BA4 and BA3 and roughly 70% of subcluster Z is composed of cells from the back of BA4. These results map out two distinct trajectories within the UMAP, from BA1 through subcluster X and from BA3-4 through subcluster Y and subcluster Z (Fig. 4G; compare the two dotted lines). Further, it is important to note that less than 6% of cells in X, Y and Z originated in BA2, suggesting these interesting subpopulations may have escaped detection within our earlier study (that analyzed smaller cell numbers) of the BA2 stream. Overall, these results present an attractive hypothesis that cells within X, Y and Z represent axial-level specific transitions between proximal neural crest cells with high SOX10 expression and distal, leading edge Trailblazers.

### Fluorescence In Situ Hybridization confirms enrichment of Trailblazer markers in neural crest cells at the distal portion of all four branchial arches

Enrichment of selected BA1-4 Trailblazer markers, including RNA Binding Motif Protein 38 (RBM38), Neurensin 1 (NRSN1), Podocalyxin-like (PODXL), and KAZALD1 was first confirmed in the cardiac mesoderm cluster (Fig. 5A-E). To visualize the in vivo expression patterns of BA1-4 Trailblazer markers, we combined fluorescent in situ hybridization (RNAscope) with immunohistochemistry for the avian neural crest marker HNK-1 (Fig. 5F). To evaluate the heterogeneity of expression of the selected markers within a typical neural crest stream, we quantified expression within individual neural crest cells, focusing on the BA2 stream (Fig. 5F). We find expression of BA1-4 Trailblazer markers within individual neural crest cells after identification of individual neural crest cells using HNK-1 (Fig. 5F). Surprisingly in mesoderm tissues distal to the neural crest cell migratory stream we also find expression of BA1-4 Trailblazer markers. For example, KAZALD1 is distinctly visible within the lead neural crest cells and in the tissue distal to the neural crest stream corresponding to lead cells within the cardiac mesoderm cluster (Suppl. Fig. 2G, 5A,C,F). NRSN1 expression was variable throughout the neural crest cell migratory stream, while expression of PODXL was enriched at the leading edge of the migratory front (Fig. 5F). We find lower expression of RBM38 within the neural crest stream, compared to the other BA1-4 Trailblazer genes selected. Thus, some Trailblazer markers confirm the position of Trailblazers at the leading edge of the migratory stream. Furthermore, the expression of some Trailblazer markers extends beyond the neural crest stream into the distil mesenchyme.

**Figure 5.**
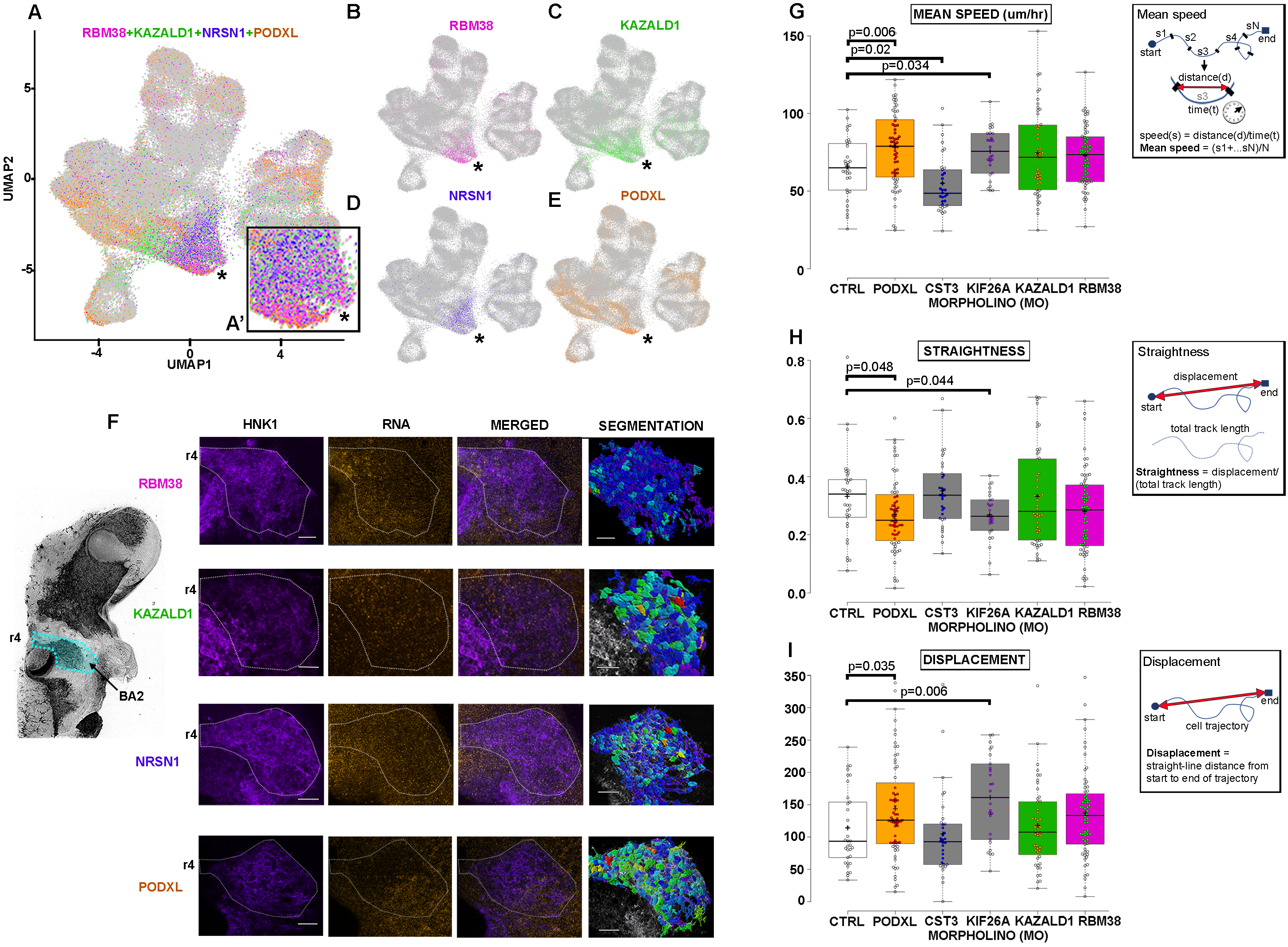
Perturbing Trailblazer markers PODXL, CST3 and KIF26A alters neural crest migration in vitro. (A-E) UMAP of BA1-4 single cell transcriptomes color-coded by expression of four genes that mark BA1-4 Trailblazers (RBM38, magenta; KAZALD1, green; NRSN1, blue; PODXL, tan). (A’) Inset is a magnification of the cardiac mesoderm cluster from (A) (asterisk). (F) Expression of individual BA1-4 Trailblazer NC cell markers by RNAscope fluorescent ISH labeling (orange) within individual BA2 NC cells labeled by IHC for HNK1 (purple). Segmentation shows each cell within a BA2 NC stream color-coded by the number of RNAscope spots for an individual BA1-4 Trailblazer marker, where blue is lowest and red is highest expression. (G-I) Knockdown of individual BA1-4 Trailblazer markers alters mean speed (G), mean speed (H), straightness (I) and displacement (I) in vitro. Branchial Arch (BA); Human Natural Killer-1 (HNK1); Rhombomere 4 (r4). Scalebar is 50um in (F).

### Perturbation of a small subset of individual Trailblazer genes altered neural crest cell behaviors in neural tube explant cultures

The knockdown of Trailblazer signature genes of BA2 led to a reduction in distance migrated by neural crest cells that colonize BA2 (Morrison 2017a). We conducted similar functional studies, to begin to assess the roles of the newly identified common BA1-4 Trailblazer signature genes within migrating neural crest cells. We knocked down a small subset (5 out of 266) of individual genes and measured changes in cell speed, straightness, and displacement from tracked cell trajectories in neural tube explant cultures (Fig. 5G-I). Morpholino knockdown of CST3 significantly decreased mean cell speed, while loss-of-function of Kinesin family member 26A (KIF26A) led to a significant decrease in cell trajectory straightness, increase in mean speed and total displacement (Fig. 5G-I). Loss-of-function of Podocalyxin-like (PODXL) led to a significant increase in cell speed and displacement, but a decrease in straightness (Fig. 5G-I). Thus, loss-of-function of a subset of BA1-4 Trailblazer signature genes (3 out of 5 tested) showed changes in neural crest cell behaviors in vitro.

### Neural crest Trailblazer genes are shared with a wide range of cell invasion phenomena in embryonic development, wound healing, and cancer metastasis

Cell invasion is a hallmark in many different biological phenomena including embryogenesis, wound repair, the immune response, and cancer metastasis. To better understand the relationship of the Trailblazer transcriptional signature with these other phenomena, we compared a gene list based on 34 published cell invasion signatures (Fig. 6A-B; Suppl. Table 4). To reduce bias, we ensured that the invasion signatures represented a broad range of cell types, techniques and model organisms. Because of the wide array of model organisms represented, we first ensured that the enriched genes curated from the 34 publications have an ortholog in chicken.

**Figure 6.**
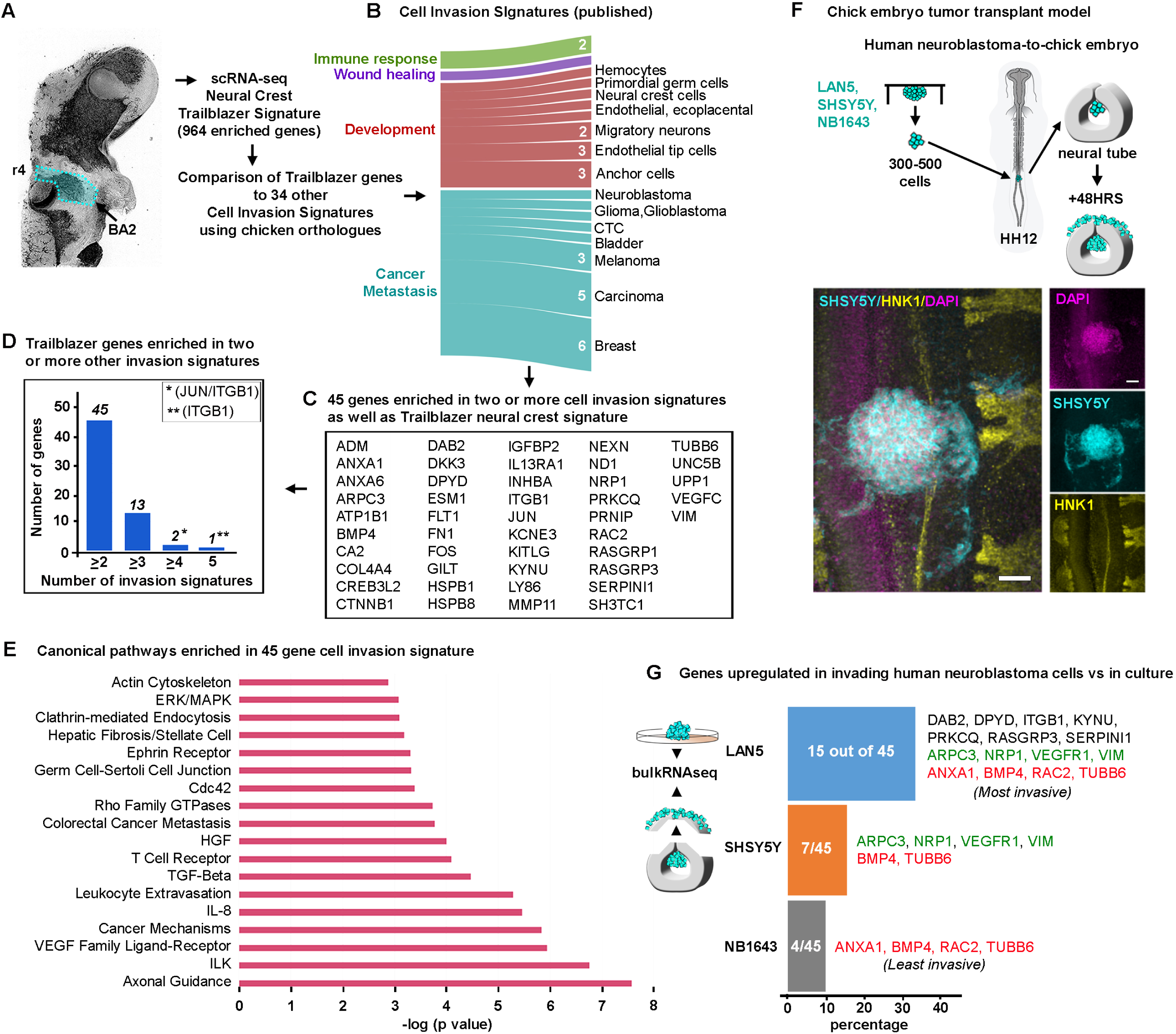
Commonalities emerge when the Trailblazer signature is compared with other invasive cell types. (A) Comparing genes enriched in BA2 Trailblazer NC cells with genes enriched in 34 invasive cell types spanning immune response, wound healing, development and metastasis (B). (C) 45 genes enriched in BA2 Trailblazers and at least 2 other invasive cell types. (D) Number genes enriched in two or more invasive cell types. (E) Canonical pathways enriched in 45 invasion signature genes. (G) Percentage of the 45 gene invasion signature enriched in migrated compared to cultured cells for neuroblastoma cell lines transplanted into the avian embryonic neural crest microenvironment. Four genes are enriched in both the least and most invasive cell types, and 2 of the 4 genes are also enriched in the SHSY5Y cells (red). Four genes are shared between the intermediate and most invasive cell types (green). 8 genes are unique to the most invasive cell type (black). Branchial Arch 2 (BA2); Rhombomere 4 (r4)

We find that 28 out of the 34 (82%) signatures displayed overlap of enriched genes with the BA2 Trailblazer signature (Morrison 2017a). There are 252 genes enriched in the Trailblazer signature that appear in at least one of the 34 published invasion signatures (Suppl. Table 4). Increasing the stringency of analysis revealed a group of 45 Trailblazer genes in 2 or more invasion signatures and 15 genes enriched in 3 or more invasion signatures (Fig. 6C,D). Two genes (JUN and ITGB1) are enriched in 4 invasion signatures and 1 gene (ITGB1) in 5 invasion signatures (Fig. 6D). We performed canonical pathway enrichment analysis and REVIGO analysis to explore specific functions and cellular states associated with the 45 gene invasion signature (Fig. 6E) (Suppl. Fig. 2I). This condensed signature, although not encapsulating all genes necessary for invasive characteristic, further supports the hypothesis that a common set of genes is required to achieve efficient collective cell invasion.

### Human neuroblastoma cells upregulate Trailblazer signature genes during invasion into the embryonic neural crest cell microenvironment

To directly test whether the 45 gene cell invasion signature determined above has direct relevance to human cancer cell invasion, we took advantage of an established chick embryo transplant model (Kulesa 2006; Bailey and Kulesa 2014). First, we transplanted 3 aggressive neural crest-derived human neuroblastoma cell lines (LAN5, SHSY5Y, NB1643) into the chick trunk embryonic neural crest microenvironment (Fig. 6F). After reincubating eggs for 48hrs, we observed that a subpopulation of neuroblastoma cells invaded the chick embryonic microenvironment along host trunk neural crest cell migratory pathways and a subset of cells remained at the transplant site in each experiment (Fig. 6F). LAN5 and SHSY5Y human neuroblastoma cells were highly invasive (Fig. 6F), but very few NB1643 cells exited from the transplant site (data not shown). To evaluate changes in gene expression, we manually isolated the chick trunk tissue containing the invasive human neuroblastoma cells and used FACs and RNA-seq analysis to compare gene expression changes to each of the cell populations in culture (Fig. 6F-G). We find that the LAN5 human neuroblastoma cells that were the most invasive in the chick embryonic neural crest microenvironment upregulated the largest percentage of the 45 gene panel, that is 15 out of 45 genes (33%) (Fig. 6C, G). Moderately invasive SHSY5Y cells upregulated a modest number of the 45 gene panel (6/45; 13%). In contrast, the poorly invasive NB1634 cells tended to remain at the transplant site, and the few invasive cells upregulated only 4 out of the 45 gene panel (less than 10%).

In comparing the profiles of the aggressive versus non-aggressive human neuroblastoma cells after transplantation into the chick embryo we noted several interesting gene expression differences (Fig. 6G). First, 6 out of 15 genes upregulated in the LAN5 cells were the same genes upregulated in the invasive SHSY5Y cells (Fig. 6G). This included actin cytoskeletal components ARPC3 (Arp2/3 related), TUBB6 (tubulin-related) and VIM (vimentin). Invasive LAN5 and SHSY5Y cells also showed upregulation of Vascular Endothelial Growth Factor (VEGF) receptors, VEGFR1 and Neuropilin-1 (NRP1), which is present in the chick embryonic microenvironment and is a neural crest cell chemoattractant (Fig. 6G, green) (McLennan 2010). NB1643 cells did not show upregulation of VEGF receptors, but included 4 genes in common with the LAN5 cells (Fig. 6G, red). These results suggest that the invasive ability of cells within the embryonic neural crest microenvironment may correlate with the number and type of genes upregulated from a common cell invasion signature.

## DISCUSSION

Cranial neural crest cells migrate in distinct streams into the branchial arches where they generate most of the bone and connective tissue of the vertebrate head. In this study we have examined the cellular composition and molecular properties of neural crest cells within the different branchial arches through the application and analysis of label free single cell transcriptional profiling of cells in the avian embryo. We find that the molecular heterogeneities within the migrating cranial neural crest of BA2 and unique transcriptional signature of the most invasive cells are a common property of cranial neural crest cells at all axial levels throughout the first four branchial arches. Furthermore, perturbation of Trailblazer genes altered neural crest cell behaviors linking them with functional roles in cell migration. We find that this Trailblazer signature is shared with a wide range of cell invasion phenomenon suggesting that the invasive ability of cells within the embryonic neural crest microenvironment correlate with genes upregulated as part of a common cell invasion signature. These findings provide a rich source of information for investigating dynamic processes in neural crest cells and raises several interesting questions with respect to neural crest biology.

Our analyses determined the cell type composition of distinct tissues within the neural crest microenvironment and identified unique gene expression differences between pre-otic and post-otic subregions as well as proximal and distal distinctions within branchial arches. We used the identification of multipotent SOX10-positive cells as a basis to discover differences in front and back cell subpopulations as well as novel, non-traditional markers of migrating neural crest cells. Distinct from SOX10-positive cells, we found a shared Trailblazer transcriptional signature within a subset of the most invasive neural crest cells within all four branchial arch streams. Trailblazer neural crest cells from all four branchial arches clustered with mesodermal cells, suggesting a high degree of transcriptional similarity between cells of two different lineages; an observation that was previously reported in zebrafish (Wagner 2018). Perturbation of a small subset of individual Trailblazer genes highly enriched within both the most invasive BA2 and BA1-4 neural crest cells (5 out of 964 and 5 out of 266 genes, respectively) revealed changes in neural crest cell behaviors in vitro. Comparison of the BA2 Trailblazer signature with other published gene expression signatures from a broad range of cell invasion phenomena revealed an overlap of 45 out of 964 genes (in 2 or more signatures). Finally, we observed that human neuroblastoma cell invasive ability correlated with the upregulation of a subset of these 45 genes in our chick embryo transplant model.

The identification of 7 distinct clusters in our UMAP analysis revealed a robust identification of cell type composition within the neural crest and first four branchial arch microenvironments, providing a knowledge base for discovery of novel cell type markers in space and time. For example, the faithful capture of HOX genes and other branchial arch specific genes to distinct subclusters within the SOX10-positive cluster provides a means for identification of novel markers within axial-level specific tissues (Suppl. Fig. 2A-F). This information also allowed us to map two distinct trajectories for the transition from SOX10-positive neural crest cells to the Trailblazer leaders (Suppl. Fig. 2H-J). Future work will be able to identify distinct spatio-temporal locations corresponding to neural crest cell hierarchy.

The surprising discovery that SOX10 and other traditional markers of migrating neural crest cells did not identify the most invasive cells suggests other novel markers must be substituted to both accurately detect the neural crest cell migratory front and entire stream. For example, the 964 genes enriched in our original Trailblazer neural crest signature were highly localized within cluster 4, rather than in cluster 1 (Fig. 3E-G) and we discovered a striking reduction of SOX10 expression in the most invasive neural crest cells (Morrison 2017a,b) (Fig. 3A-D; Suppl. Fig. 2G). Thus, traditional markers of migrating neural crest cells used to mark cell position and/or specifically isolate neural crest cells may not fully capture the most invasive neural crest cells. This may lead to mistakes in assessing the timing and position of the neural crest cell streams and incomplete gene expression analysis of migrating neural crest cells common in typical isolation-required RNA-seq methods. We suggest that COL18A1, whose expression was previously described at the enteric neural crest cell wavefront (Nagy 2018), more faithfully captures migrating neural crest cells at both the front and back of the migratory streams (Suppl. Fig. 2H) an may offer a possible solution.

In vivo single cell analysis by multiplexed FISH confirmed a repertoire, rather than enhanced expression of any individual gene, represents the most invasive neural crest cells. Expression analysis of a small subset of BA1-4 Trailblazer genes confirmed the heterogeneous expression within the lead neural crest cells that we discovered in our previous extensive validation of Trailblazer gene expression (Morrison 2017a). This suggested that gain- or loss-of-function of any individual Trailblazer gene may only have modest effects on cell behaviors. This was indeed observed in vitro after loss-of-function of a small subset of individual Trailblazer genes in neural tube explant cultures, with only loss-of-function of PODXL, CST3, or KIF26A showing significant changes in cell speed, straightness, or displacement (Fig. 5). This highlighted the functional redundancy and robustness of neural crest cell migration that underlies these complex cell behaviors as critical to the formation of craniofacial and cardiovascular development. Further, neural crest cell-cell communication may overcome minor changes to a subset of cells since our knockdown method only affected a subset of the population of all premigratory neural crest. Future technical methods that allow for multiple gene manipulation in a systematic manner and knockdown in every cell either in silico, in vitro, or in vivo may help to shed light on the Trailblazer network dynamics of neural crest cell invasion.

A subset of Trailblazer neural crest genes was shared in a broad range of cell invasion phenomena suggesting common collective cell migration mechanisms. In our attempt to be inclusive, we used molecular signatures that were acquired from a wide variety of methods, model organisms, and cell lines (Fig. 6). Importantly, not all cell invasion signatures included a complete transcriptome analysis. Another limitation of this comparison was a lack of exact orthologues between species. Nevertheless, the overlap of 45 out of 964 genes does provide unique molecular inroads and the in vivo neural crest model is poised to study gene network dynamics and function. This is exemplified by our results that showed human neural crest-derived neuroblastoma cells transplanted into our chick embryo model upregulated Trailblazer genes according to invasive ability and offer a rapid means to test gene candidates in human cancer metastasis.

In summary, our findings offer unprecedented single cell resolution of the molecular expression patterns that underlie neural crest cell migration and formation of the first four branchial arches during vertebrate development. Previous knowledge derived from our single cell qPCR and RNA-seq profiling of purified neural crest cells combined with the innovative label-free, non-isolation cell profiling of the entire branchial arches provided confidence in mapping cell type composition and a much richer data set to examine both neural crest and neural crest-microenvironment signaling. This allowed us to identify axial level and spatio-temporal molecular heterogeneities within the first four branchial arches; insights that would have been lost in a bulk RNA-seq analysis of the tissues or purified neural crest cells. These insights included novel markers of pre-otic, post-otic and front-to-back cell populations and the discovery of a refined Trailblazer transcriptional signature of the most invasive neural crest cells within all four branchial arch streams. Mapping of the genes enriched in this refined signature to specific signaling pathways will provide for future in-depth functional experiments that begin to tease out the complex network dynamics that underlie collective cell migration and cellular hierarchy of the neural crest. The molecular signature of the neural crest compared well with other cell invasion phenomena, suggesting the exciting possibility that fundamental properties underlie development, wound healing, and cancer metastasis. The neural crest model combined with our dynamic in vivo imaging platform present a clear means to address complex questions in the biology of collective cell migration.

## METHODS

### Single cell isolation of chick tissue

All experiments were performed according to institutional and federal ethical standards. Fertilized, white leghorn chicken eggs (NCBI Taxonomy ID:9031; Centurion Poultry, Lexington, GA, USA) were incubated at 38 degrees C in a humidified incubator until the desired Hamburger and Hamilton (HH) stage (Hamburger and Hamilton 1951) of development. Embryos were screened for health and harvested into chilled 0.1% DEPC phosphate-buffered saline (PBS). To capture neural crest during active migration, branchial arches (BA) 1 and 2 were manually isolated from HH13 embryos (n=15 embryos). Branchial arches 3 and 4 were manually isolated from HH15 embryos (n=10 embryos). Each branchial arch was further manually dissected into the front 20% (distal-most) and 80% back (proximal-most) portions. Stage-, branchial arch- and tissue portion-matched tissues were pooled and dissociated as previously described (Morrison 2015). The viability and concentration of each single cell suspension was quickly confirmed on a Nexelome Cellometer Auto T4 (Nexelome Bioscience, Lawrence, MA, USA) and the cell suspensions used as input for 10X scRNA-seq (10X Genomics, San Francisco, CA, USA) per the manufacturer’s recommendations.

### 10x Chromium single-cell RNA-seq library construction

Dissociated cells were loaded on a Chromium Single Cell Controller (10x Genomics, Pleasanton, CA, USA), based on live cell concentration, with a target of 4,000-10,000 cells per sample. Libraries were prepared using the Chromium Single Cell 3’ Library & Gel Bead Kit v2 (10x Genomics, Pleasanton, CA, USA) according to manufacturer’s directions. Resulting short fragment libraries were checked for quality and quantity using an Bioanalyzer 2100 (Agilent Technologies, Santa, Calara, CA, USA) and Qubit Fluorometer (Invitrogen, Waltham, MA, USA). Libraries were pooled and sequenced to a depth necessary to achieve 25-35,000 mean reads per cell, 125-520M reads each, on an Illumina HiSeq 2500 (Illumina, San Diego, CA, USA) instrument using Rapid SBS v2 chemistry with the following paired read lengths: 26 bp Read1, 8 bp I7 Index and 98 bp Read2.

### Bioinformatics Analysis

Eight samples of 10x Genomics Chromium scRNA-seq data were sequenced on five Illumina HiSeq 2500 flowcells (Illumina, San Diego, CA, USA). Data was processed with bcl2fastq (2.20) and aligned and aggregated with CellRanger (2.1.0). Data was aligned to galGal4 using annotations from Ensembl 84. Downstream analysis was done in R (3.5.0) with the Seurat package (2.3.4) (Butler 2018). Cells were kept for downstream analysis if they had more than 500 genes expressed, less than 20,000 UMIs, and less than 30% mitochondrial expression. 2796 genes were selected as variable genes for downstream analysis, and the first 50 principal components were used for later steps (UMAP and cluster identification). For identifying cluster markers, we used FindAllMarkers with the minimum percentage of cells within a cluster expressing the gene set to 25% and a log fold change threshold of greater than or equal to 0.25. To further investigate some clusters, we performed subclustering by subsetting the Seurat object by cluster and running FindClusters at higher resolutions. UMAP has gained favor in recent single cell analyses because it retains more of the global architecture of the data set (McInnes 2018; Becht 2019) and information may be gathered from the spatial relationship of single cell clusters (Becht 2019).

### Fluorescent In Situ Hybridization by RNAscope and Immunohistochemistry

Integrated analysis of gene expression and protein detection in single cells was performed as previously described (Morrison 2017a,b). Briefly, RNAscope (Advanced Cell Diagnostics, Newark, CA, USA) fluorescent in situ hybridization and subsequent immunohistochemistry were carried out on fixed avian embryos. Images were acquired on a Zeiss LSM 800 laser scanning confocal microscope (Carl Zeiss, Jena, Germany) and analyzed in Imaris (Bitplane, Belfast, Northern Ireland). After the 3D boundaries of each neural crest cell were determined by the membrane-specific HNK-1 IHC signal, the number of RNAscope spots within the 3D volume of each neural crest cell were quantified.

### In vitro neural crest cell cultures

In vitro neural tubes cultures and cell tracking analysis were performed as previously described (McLennan 2010; McLennan 2020). 2 hours prior to neural tube removal from the embryos, the dorsal portion of each neural tube was transfected with fluorescein-tagged splice modifying morpholinos targeting the trailblazer genes using in ovo electroporation as previously described (McLennan and Kulesa 2019). The 5’ to 3’ morpholino sequences are CGATTTTAATCTTCTGATACCTGCT for PODXL, ACATCCTGCTCAGAGCCTACCTTAG for CST3, AACAGAAAGGTGACAAACCTGATGA for Kif26a, ATCTGAGGCTCTGGGAATGGAAGAT for KAZALD1, GACGGACCTCTAAGGCTCCTCACC for RBM38 (Gene Tools, Philomath, OR, USA).

### Neuroblastoma Cell Lines

Human neuroblastoma cell lines LAN5, NB1643 (Children’s Oncology Group, Texas Tech University, USA) and SHSY5Y (ATCC, Manassas, VA, USA; CRL-2266) were maintained as in Kasemeier-Kulesa et at. 2018. Neuroblastoma cells were cultured in hanging drops and transplanted into Hamburger Hamilton (HH) stage 10 chick embryos (Kulesa 2006).

## ACKNOWLEDGEMENTS

PMK would like to acknowledge the kind and generous funding from the Stowers Institute for Medical Research.

## DATA AVAILABILITY

Original data underlying this manuscript can be accessed from the Stowers Original Data Repository at https://www.stowers.org/research/publications/odr. All raw and processed sequencing data generated in this study have been submitted to the NCBI Gene Expression Omnibus (GEO; https://www.ncbi.nlm.nih.gov/geo/) under accession number GSEXXXXXX.

